# Knowledge, Attitudes and Practices on Antimicrobial Resistance Among Practising Veterinary and Para-veterinary Professionals in Zambia

**DOI:** 10.1101/2025.08.14.670270

**Authors:** Trascilla Mwape, Tamsin Dewé, Doreen Sitali, Luke Nyakarahuka, Claire Gilbert, Alison Pyatt, John Bwalya Muma, Chisoni Mumba

## Abstract

In this study, we aimed to assess the knowledge, attitudes, and practices (KAP) of veterinary and para-veterinary professionals concerning antimicrobial resistance (AMR), in order to identify key drivers of inappropriate antimicrobial use and gaps in current stewardship efforts. Data was collected from a cross-sectional survey of 144 veterinary and para-veterinary professionals (n=144) using an online structured self-administered questionnaire in ZOHO software. We analyzed data using descriptive statistics and logistic regression in IBM SPSS Statistics Version 25.

The findings revealed a high foundational knowledge of AMR, with 66.7% correctly defining AMR. The majority of respondents (82%) acknowledged that overuse of antimicrobials reduces their effectiveness and expressed strong support for mitigation efforts, with 95.8% endorsing farmer education as a key strategy. However, significant gaps in practice persisted: 75% of animal owners frequently initiated antibiotic use without supervision; professionals largely relied on empirical treatment (62.5%); and only 20.8% of professionals routinely performed antimicrobial susceptibility testing (AST), due to barriers like cost and laboratory accessibility. Logistic regression analysis revealed no significant associations between socio-demographic variables and levels of AMR knowledge or attitudes. This suggests that barriers to responsible antimicrobial use may be more structural than individual.

We conclude that while Zambian practicing veterinary and para-veterinary professionals possess strong AMR awareness and positive attitudes, risky practices do persist. We recommend strengthening AMR stewardship with a holistic One Health approach by improving veterinary services to promote more appropriate and effective antimicrobial use.

**Author Summary:** Antibiotics are essential for treating sick animals, but their misuse can lead to antimicrobial resistance, where bacteria no longer respond to the drugs meant to kill them. This problem threatens both animal and human health. In Zambia, veterinary and para-veterinary professionals are actively involved in administering treatments and supporting decisions related to antibiotic use in livestock and pets. We wanted to understand what these professionals know, believe, and do when it comes to using antibiotics responsibly.

To find out, we surveyed 144 veterinary and para-veterinary professionals from across the country. We learned that most of them were aware that misuse of antibiotics can make treatments less effective and that drug-resistant bacteria can spread between animals and people. Many also supported educating farmers and working with human health experts to tackle the issue. However, we also discovered that antibiotics were often used in animals without proper testing or diagnosis, mainly because of limited laboratory access and cost concerns.

Our findings show that while knowledge and attitudes are strong, actual practices need improvement. We believe improved access to diagnostic services, more explicit rules, and ongoing training can help veterinary and veterinary paraprofessionals protect both animal and public health from the growing threat of antibiotic resistance.

## 1.0 Introduction

Antimicrobial resistance (AMR) constitutes a significant global threat to both human and animal health, primarily due to the misuse and overuse of antimicrobials (1). This phenomenon involves microorganisms developing the ability to thrive in the presence of drugs intended to inhibit their growth or eliminate them (2). Resistant bacteria can also be transmitted between animals, people, food, and the environment. The complexity of AMR extends beyond endangering human and animal well-being, posing substantial risks to regional, national, and global security, as well as economic stability (3).

In the Zambian context, where agriculture and animal husbandry are pivotal to the economy, the responsible use of antimicrobial agents is of paramount importance (4). Veterinary and para-veterinary professionals, being at the forefront of managing animal health, play a crucial role, and their knowledge and practices directly influence the emergence and spread of antimicrobial resistance. The responsible use of antimicrobial drugs in veterinary medicine is essential to mitigate the emergence and spread of resistance (5).

The primary drivers of AMR are often attributed to the widespread utilization of antimicrobials across various sectors (6). This contributes significantly to the dissemination of resistant pathogens and resistance elements within and among these sectors (7). Factors influencing the use of antimicrobials include knowledge, expectations, the nature of practices, interactions between prescribers and patients, economic incentives, health system characteristics, and the regulatory environment (8).

To address the escalating issue of AMR, international organisations such as the World Health Organisation (WHO), the World Organisation for Animal Health (OIE), and the Food and Agriculture Organisation (FAO) have devised strategies (9). Particularly noteworthy is the FAO’s AMR action plan, recognizing the intricate links between human, animal, and environmental health (10). This plan concentrates on five key objectives: enhancing awareness and understanding, fortifying AMR-related governance, advocating best practices in animal production and health, enhancing surveillance and monitoring, and fostering research and innovation (10). More recently, the 2024 political declaration of the High-level meeting on AMR has reinforced these priorities and gone further by setting measurable global targets, including a 10% reduction in AMR-related deaths by 2030 (11). The declaration also calls for an independent scientific panel to guide AMR policy and emphasises equitable access to antimicrobials, especially in low- and middle-income countries (11). This builds on earlier efforts like those of the FAO, which focused more on sector-specific interventions and capacity development within food and agriculture systems. Veterinary professionals play a crucial role in implementing these objectives and promoting responsible antimicrobial use in animal health (12).

In the absence of prior research on AMR knowledge, attitudes, and practices among veterinary and para-veterinary professionals in Zambia, this study provides essential baseline insights into this critical area. Veterinary and para-veterinary professionals play a crucial role in administering antimicrobials to animals. Given the livestock sector’s key role in Zambia’s economy, the misuse of antimicrobials in this sector increases the risk of AMR development and dissemination, which can, in turn, affect public health. Thus, evaluating the knowledge of veterinary and para-veterinary professionals is essential to promote responsible antimicrobial use and safeguard animal and public health.

## 2.0 Methodology

### Ethics statement

We obtained ethical clearance for this study from the University of Zambia Biomedical Research Ethics Committee (UNZABREC) and obtained administrative permission from the National Health Research Authority under reference number **NHRA-1311/20/06/2024**. The first page of the structured online questionnaire in ZOHO software was a consent form, and participants were required to consent before completing the questionnaire. We communicated that participation was voluntary and that individuals could withdraw from the study at any point without facing any consequences. We anonymized all collected data to protect participant confidentiality, and only members of the research team had access to the information. We also assured participants that their responses would remain confidential and would not impact their professional standing in any way.

### 2.1 Study design

We employed a cross-sectional study design to assess the knowledge, attitudes, and practices related to antimicrobial resistance (AMR) among practicing veterinary and para-veterinary professionals in Zambia. Our study population included both public and private sector practicing professionals actively working in livestock/large animal, small animal/companion animal, and poultry sectors.

We defined a veterinary professional as a practicing registered veterinarian who completed a six-year Bachelor of Veterinary Medicine (BVM) program and received certification from the Veterinary Council of Zambia (VCZ). Similarly, we considered a practicing registered para-veterinarian to be a trained professional holding a Certificate, Diploma, or Degree in animal science, who assists veterinarians with fieldwork and has also been certified by the VCZ. The inclusion criteria were veterinarians and para-veterinary professionals who have been practicing for at least two years, as well as those actively engaged in their respective fields at the time of the study. Similarly, we excluded veterinarians and para-veterinarians who have retired from active clinical practice, and those practicing outside Zambia.

### 2.3 Description of the Study Sites

Veterinarians and para-veterinarians are widely distributed in Zambia, with each of the 116 districts (13) being run by one veterinary officer and several veterinary camps manned by veterinary assistants (para-veterinary professionals). We adopted a national approach in this study, ensuring participation from all ten provinces of Zambia as show in Figure 1. The majority of responses came from Lusaka Province (35.39%) and Southern Province (33.15%), reflecting their relatively high concentration of veterinary and para-veterinary professionals. Other provinces also contributed to the study, including Western (7.30%), Central and Northern (5.06% each), Northwestern (4.49%), Copperbelt (3.93%), Eastern (2.81%), Muchinga (1.69%), and Luapula (1.12%) as shown in Figure 1. This distribution highlights the nationwide scope of our data collection and reinforces the representativeness of our findings across Zambia.

**Fig 1.**
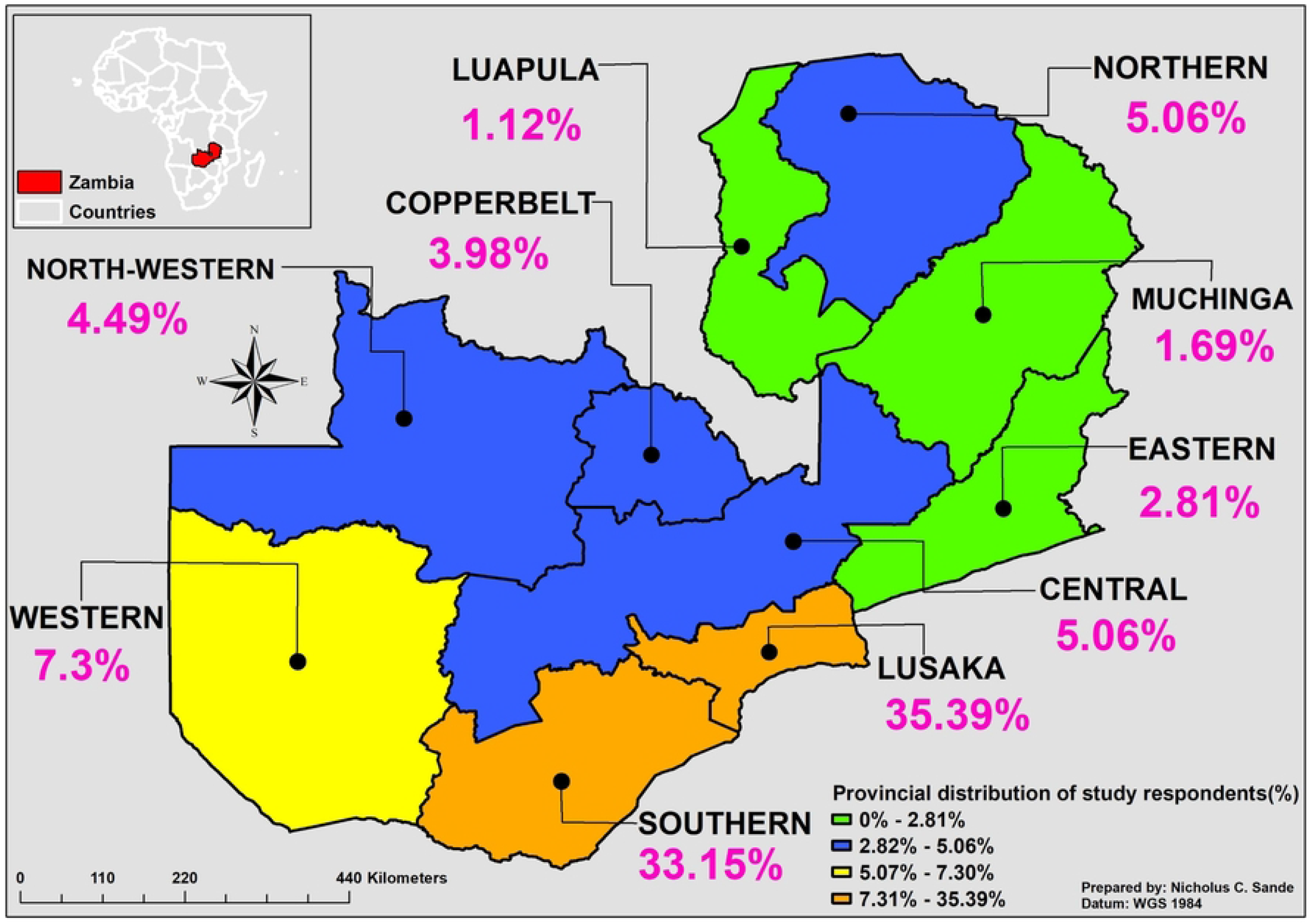
Map of Zambia showing study sites and proportion of the sample size. Developed by the authors. Reprinted from [Zambia_Mosaic_250Karc1950_ddecw] under a CC BY licence, with permission from the Surveyor General, Government Republic of Zambia, original copyright, [1974].

### 2.4 Sample size estimation

Zambia has a total of 441 veterinarians and 1358 para-veterinary professionals registered with the Veterinary Council of Zambia (VCZ). The Ministry of Fisheries and Livestock is the biggest employer of veterinarians and para-veterinary professionals. Since we did not have the exact number of practising veterinarians and paraprofessionals, we calculated a minimum number of 116 respondents taking a national approach. With a non-response rate of 10%, the sample size was adjusted to 127 veterinary and para-veterinary professionals. To reach our target population, we administered an online structured questionnaire through professional social media groups commonly used by veterinary and para-veterinary professionals. As anticipated, Southern and Lusaka provinces recorded the highest number of practicing veterinarians and para-veterinary professionals. This aligns with regional practice patterns, with Lusaka serving as the capital city and commercial hub, where a large urban population supports a higher concentration of small animal practitioners (14). In comparison, Southern Province has extensive cattle-rearing and livestock production systems, which explains the predominance of large animal and livestock practitioners in the region (14).

### 2.5 Data collection techniques

A structured questionnaire was developed to collect data on demographics, knowledge, attitudes, practices related to antibiotic use and resistance, and awareness of antimicrobial stewardship. We pretested the questionnaire among 12 veterinarians and para-veterinarians in Lusaka District. Based on their feedback, we refined the questions to enhance clarity and relevance. We excluded those who participated in the pretest from the final survey.

The finalized questionnaire was self-administered virtually to participants across the country. Data collection took place between August and December 2024.

### 2.6 Data management and analysis

We first recorded the collected data in Microsoft Excel for coding and cleaning. During this process, we excluded entries with missing information. After cleaning, we transferred the data to the IBM Statistical Package for Social Sciences (SPSS Version 25) software for statistical analysis.

We conducted descriptive analysis using frequencies and percentages to summarize participant responses. To assess knowledge, attitudes, and practices, we used a five-point Likert scale, which was rated as follows: Strongly Agree (SA) corresponding to a score of 5, Agree (A) to 4, Neutral (N) to 3, Disagree (D) to 2, and Strongly Disagree (SD) to 1. The scoring direction (e.g., 5 for correct, 1 for incorrect) was reversed as appropriate depending on the nature of the question to ensure consistency in interpretation. Each participant’s assessment was based on ten questions, totaling a maximum of 50 points. The aggregated scores were then converted into percentage form and categorized as either “good” or “poor” using Bloom’s cut-off point, with a score ranging from 80-100% being classified as good, 60-<80% being classified as moderate, and below 60% being rated as poor. This identical scoring criterion was consistently applied to questions assessing both knowledge and attitude.

For practices, we used descriptive statistics to identify common trends in antimicrobial use. We reported these trends using frequencies and percentages, focusing on guideline adherence, prescribing behaviours, and notable variations among respondents. Unlike knowledge and attitude, we did not categorize practices into performance groups; instead, we grouped them into “risky” and “non-risky” categories.

After descriptive analysis, logistic regression was performed to assess the associations between the binary outcome variables (categorized as “good” versus “poor” knowledge, and “good” versus “poor” attitude) of the respondents concerning antibiotic usage and AMR. The independent variables included in this analysis were gender, age, level of qualification, type of practice, and work experience of the respondents. These specific independent variables were carefully selected based on their theoretical relevance and potential influence on knowledge and attitude towards antibiotic usage and AMR. Categorical variables such as gender and type of practice were appropriately dummy coded. Age and work experience were treated as continuous variables, while the level of qualification was categorized into distinct levels (e.g., Bachelor’s degree, Master’s degree, etc.) to reflect varying educational backgrounds. The fit of the logistic regression model was rigorously assessed using the likelihood ratio test. The strength and direction of the associations were subsequently determined through the calculation of odds ratios (AOR) and their corresponding 95% confidence intervals, with statistical significance set at p<0.05.

## 3.0 RESULTS

A total of 144 veterinary and para-veterinary professionals participated in the study (Table 1). The majority were female (63.2%), with males accounting for 36.8%. The participants were predominantly young to mid-career professionals, with over 63% falling within the 26 to 40-year age range. Notably, no respondents were below the age of 25, and only a small proportion (15.2%) were above 50 years.

**Table 1:** Summary of socio-demographic information of veterinary and para-veterinary professionals.

| Variable | Category | Frequency | Proportion (%) |
| --- | --- | --- | --- |
| Sex | Female | 91 | 63.2 |
|  | Male | 53 | 36.8 |
| Age | < 25 | 0 | 0.0 |
|  | 26 – 30 | 31 | 21.5 |
|  | 31 – 35 | 28 | 19.4 |
|  | 36 – 40 | 32 | 22.2 |
|  | 41 – 45 | 14 | 9.7 |
|  | 46 – 50 | 17 | 11.8 |
|  | 51 – 55 | 11 | 7.6 |
|  | >55 | 11 | 7.6 |
| Marital status | Married | 110 | 76.4 |
|  | Single/divorced | 34 | 23.6 |
| Education | Certificate | 30 | 20.8 |
|  | Diploma | 20 | 13.9 |
|  | Degree | 55 | 38.2 |
|  | Masters | 32 | 22.2 |
|  | PhD | 7 | 4.9 |
| Veterinary Training | UNZA | 88 | 61.1 |
|  | ZIAH | 45 | 31.3 |
|  | Abroad | 7 | 4.9 |
|  | NRDC | 3 | 2.1 |
|  | MU | 1 | 0.7 |
| Occupation | Government | 94 | 65.3 |
|  | Private | 41 | 28.5 |
|  | Researcher/Teacher | 9 | 6.2 |
| Years of practice | 2 – 5 | 38 | 26.4 |
|  | 6 – 10 | 31 | 21.5 |
|  | 11 – 15 | 35 | 24.3 |
|  | 16 – 20 | 13 | 9.0 |
|  | 21 – 25 | 11 | 7.6 |
|  | >25 | 16 | 11.1 |
| Type of veterinary practice | Mixed | 110 | 76.4 |
|  | Poultry | 8 | 5.6 |
|  | Small | 6 | 4.2 |
|  | Large | 5 | 3.5 |
|  | Aquatic | 5 | 3.5 |
|  | Exotics | 3 | 2.1 |
|  | Wildlife | 7 | 4.9 |

Regarding marital status, 76.4% were married, while 23.6% were single or divorced. In terms of educational attainment, over a third (38.2%) of participants held a degree, followed by those with master’s (22.2%) and certificate qualifications (20.8%). A smaller proportion had attained a PhD (4.9%).

The majority received their veterinary training from the University of Zambia (61.1%), with others having trained at the Zambia Institute of Animal Health (ZIAH) (31.3%), abroad (4.9%), or at other institutions such as the Natural Resource Development College (NRDC) and Mulungushi University (MU).

Occupationally, most respondents were employed in the government sector (65.3%), while 28.5% worked in the private sector, and 6.2% were researchers or educators. Experience levels varied, with 72.2% having between 2 and 15 years of practice, indicating a relatively young workforce.

The most common type of veterinary practice was mixed practice (76.4%), suggesting broad involvement across various species. Specialized practices such as poultry, wildlife, aquatic, and exotic animal care were less common (Table 1).

### 3.2 Knowledge of AMR

The results indicate that 5.6% of respondents had low knowledge of AMR concepts, highlighting the need for targeted education and capacity building (Table 2). A further 7.6% demonstrated moderate understanding, pointing to an opportunity for strengthening foundational knowledge through continuous professional development. Encouragingly, 86.8% achieved high scores, suggesting the presence of well-informed professionals who could act as AMR champions or peer educators as shown in Table 2.

**Table 2:** Knowledge regarding antimicrobial resistance among veterinary and para-veterinary professionals.

| Knowledge variables | Category | Frequency n=144 | Proportion (%) |
| --- | --- | --- | --- |
| AMR is a microorganism's ability to withstand drugs meant to kill or inhibit it | Strongly disagree | 7 | 4.86 |
|  | Disagree | 1 | 0.69 |
|  | Neutral | 2 | 1.39 |
|  | Agree | 38 | 26.39 |
|  | Strongly agree | 96 | 66.67 |
| Resistant bacteria can spread between animals and humans | Strongly disagree | 6 | 4.17 |
|  | Disagree | 0 | 0.0 |
|  | Neutral | 9 | 6.25 |
|  | Agree | 57 | 39.58 |
|  | Strongly agree | 72 | 50.00 |
| Antibiotics kill both commensal and pathogenic bacteria | Strongly disagree | 7 | 4.86 |
|  | Disagree | 11 | 7.64 |
|  | Neutral | 16 | 11.11 |
|  | Agree | 60 | 41.67 |
|  | Strongly agree | 50 | 34.72 |
| Overuse of antibiotics renders them ineffective | Strongly disagree | 7 | 4.86 |
|  | Disagree | 10 | 6.94 |
|  | Neutral | 8 | 5.56 |
|  | Agree | 58 | 40.28 |
|  | Strongly agree | 61 | 42.36 |
| Microorganisms can become resistant to antibiotics | Strongly disagree | 6 | 4.17 |
|  | Disagree | 1 | 0.69 |
|  | Neutral | 0 | 0.0 |
|  | Agree | 72 | 0.50 |
|  | Strongly agree | 65 | 45.14 |
| It is important to observe withdrawal periods for milk and meat after antibiotic treatment | Strongly disagree | 10 | 6.94 |
|  | Disagree | 0 | 0.00 |
|  | Neutral | 0 | 0.00 |
|  | Agree | 21 | 14.58 |
|  | Strongly agree | 113 | 78.47 |
| It is important to administer antibiotics with the correct dose and dosage for all animal species | Strongly disagree | 9 | 6.25 |
|  | Disagree | 0 | 0.00 |
|  | Neutral | 0 | 0.00 |
|  | Agree | 21 | 14.58 |
|  | Strongly agree | 114 | 79.17 |
| Treating the entire herd with antibiotics can prevent infection spread | Strongly disagree | 62 | 43.06 |
|  | Disagree | 50 | 34.72 |
|  | Neutral | 11 | 7.64 |
|  | Agree | 6 | 4.17 |
|  | Strongly agree | 15 | 10.42 |
| Antibiotics for healthy animals can prevent future illness | Strongly disagree | 52 | 36.11 |
|  | Disagree | 61 | 42.36 |
|  | Neutral | 7 | 4.86 |
|  | Agree | 17 | 11.80 |
|  | Strongly agree | 7 | 4.86 |
| Infections are becoming increasingly resistant to treatment using antibiotics | Strongly disagree | 5 | 3.47 |
|  | Disagree | 3 | 2.08 |
|  | Neutral | 7 | 4.86 |
|  | Agree | 90 | 62.50 |
|  | Strongly agree | 39 | 27.10 |

### 3.3 Attitude towards AMR

The results indicate that a majority of veterinary and para-veterinary professionals recognize the significance of antimicrobial resistance in their field. For instance, 92% of the respondents agreed that veterinary professionals play a crucial role in combating AMR, and 93.8% acknowledged AMR as a growing concern in Zambia’s livestock sector (Table 3). Similarly, 95.8% agreed that farmers need education on antimicrobial alternatives, and 96.5% emphasised the importance of AMR training and workshops for veterinary and para-veterinary professionals. Attitudes toward restricting antimicrobial access were also generally positive, with 87.5% agreeing that restricting veterinary antimicrobial access helps combat AMR. Overall, while 75% of the respondents maintained moderate to high levels of positive attitudes toward AMR issues, approximately 25% still hold suboptimal attitudes, highlighting the need for targeted educational interventions as shown in Table 3.

**Table 3:** Attitude towards AMR among Veterinary and para-veterinary professionals.

| Attitude variables | Category | Frequency n=144 | Proportion (%) |
| --- | --- | --- | --- |
| Veterinary and para-veterinary professionals help combat AMR | Strongly disagree | 4 | 2.8 |
|  | Disagree | 3 | 2.1 |
|  | Neutral | 4 | 2.8 |
|  | Agree | 38 | 26.4 |
|  | Strongly agree | 95 | 66.0 |
| AMR is a growing concern in Zambia's livestock sector | Strongly disagree | 4 | 2.8 |
|  | Disagree | 2 | 1.4 |
|  | Neutral | 3 | 2.1 |
|  | Agree | 49 | 34.0 |
|  | Strongly agree | 86 | 59.7 |
| AMR is a significant public health problem in Zambia | Strongly disagree | 4 | 2.8 |
|  | Disagree | 1 | 0.7 |
|  | Neutral | 10 | 6.9 |
|  | Agree | 46 | 31.9 |
|  | Strongly agree | 83 | 57.6 |
| Farmers need education on antimicrobial alternatives in animal care | Strongly disagree | 5 | 3.5 |
|  | Disagree | 0 | 0.0 |
|  | Neutral | 1 | 0.7 |
|  | Agree | 46 | 31.9 |
|  | Strongly agree | 92 | 63.9 |
| Restricting veterinary antimicrobial access helps combat AMR | Strongly disagree | 8 | 5.6 |
|  | Disagree | 4 | 2.8 |
|  | Neutral | 6 | 4.2 |
|  | Agree | 48 | 33.3 |
|  | Strongly agree | 78 | 54.2 |
| AMR training and workshops are vital for veterinary professionals | Strongly disagree | 2 | 1.4 |
|  | Disagree | 1 | 0.7 |
|  | Neutral | 2 | 1.4 |
|  | Agree | 47 | 32.6 |
|  | Strongly agree | 92 | 63.9 |
| Routine AMR surveillance in veterinary pathogens guides treatment | Strongly disagree | 3 | 2.1 |
|  | Disagree | 0 | 0.0 |
|  | Neutral | 2 | 1.4 |
|  | Agree | 58 | 40.3 |
|  | Strongly agree | 81 | 56.3 |
| Collaboration with human health professionals is key to combating AMR | Strongly disagree | 3 | 2.1 |
|  | Disagree | 0 | 0.0 |
|  | Neutral | 1 | 0.7 |
|  | Agree | 39 | 27.1 |
|  | Strongly agree | 101 | 70.1 |
| Thorough animal examination is essential before prescribing antibiotics | Strongly disagree | 3 | 2.1 |
|  | Disagree | 1 | 0.7 |
|  | Neutral | 1 | 0.7 |
|  | Agree | 80 | 55.6 |
|  | Strongly agree | 59 | 41.0 |
| Advanced diagnostics are essential for accurate pathogen identification and treatment | Strongly disagree | 2 | 1.4 |
|  | Disagree | 0 | 0.0 |
|  | Neutral | 3 | 2.1 |
|  | Agree | 57 | 39.6 |
|  | Strongly agree | 82 | 56.9 |

### 3.4 Practices related to antimicrobial resistance

These results indicate a moderate to poor level of AMR stewardship practices among veterinary and para-veterinary professionals in Zambia. A high percentage (75%) reported that animal owners frequently start antibiotics without veterinary input, and 62.5% use empirical treatment for first-time patients (Table 4). AST is rarely done, with 73.61% admitting they seldom or never perform it, and 49.30% prescribe antimicrobials without thorough examination. Collaboration with AMR specialists is also low (62.50%), and key barriers to AST include lack of lab services (57.64%), cost (24.31%), urgency (11.11%), and long waiting times (16.67%). However, some positive practices were noted. About 52.08% request AST after poor treatment outcomes, and 21.53% do so for recurrent cases. Most rely on formal training (69.44%) and scientific literature (12.50%) for antibiotic information as shown in Table 4.

**Table 4:** Risky and non-risky practices among Veterinary and para-veterinary professionals in Zambia.

| Practice Area | Specific Behaviour | Risk Classification | Proportion (%) |
| --- | --- | --- | --- |
| Animal owners initiating antibiotics without veterinary supervision | Frequently | Risky | 75.00 |
| Choice of antibiotic for a first-time patient | Empirical treatment | Risky | 62.50 |
| Performance of AST before prescribing | Rarely or never | Risky | 73.61 |
| Prescribing antimicrobials without a thorough examination | Frequently or occasionally | Risky | 49.30 |
| Collaboration with AMR specialists | Rarely or never | Risky | 62.50 |
| Barriers to AST – Lack of lab services | Reported as a barrier | Risky | 57.64 |
| Barriers to AST – Inability of owners to pay | Reported as a barrier | Risky | 24.31 |
| Barriers to AST – Urgent need for antibiotics | Reported as a barrier | Risky | 11.11 |
| Barriers to AST – Long waiting time | Reported as a barrier | Risky | 16.67 |
| Reason for requesting AST | Poor response to previous treatment | Non-Risky | 52.08 |
| Reason for requesting AST | Recurrent health conditions | Non-Risky | 21.53 |
| Source of antibiotic information | Veterinary education/training | Non-Risky | 69.44 |
| Source of antibiotic information | Scientific literature | Non-Risky | 12.50 |
| Encountering AMR cases in practice | Frequently or occasionally | Non-Risky | 68.06 |
| Follow-up on antimicrobial treatment effectiveness | Frequently or occasionally | Non-Risky | 84.72 |
| Participation in AMR awareness campaigns | Frequently or occasionally | Non-Risky | 59.72 |

### 3.5 Determinants of AMR

To assess predictors of knowledge and attitudes, logistic regression analysis was conducted. The results show that none of the variables under consideration were significant predictors of knowledge and attitudes regarding AMR (AOR > 0.05) as shown in Tables 5 and 6.

**Table 5:**
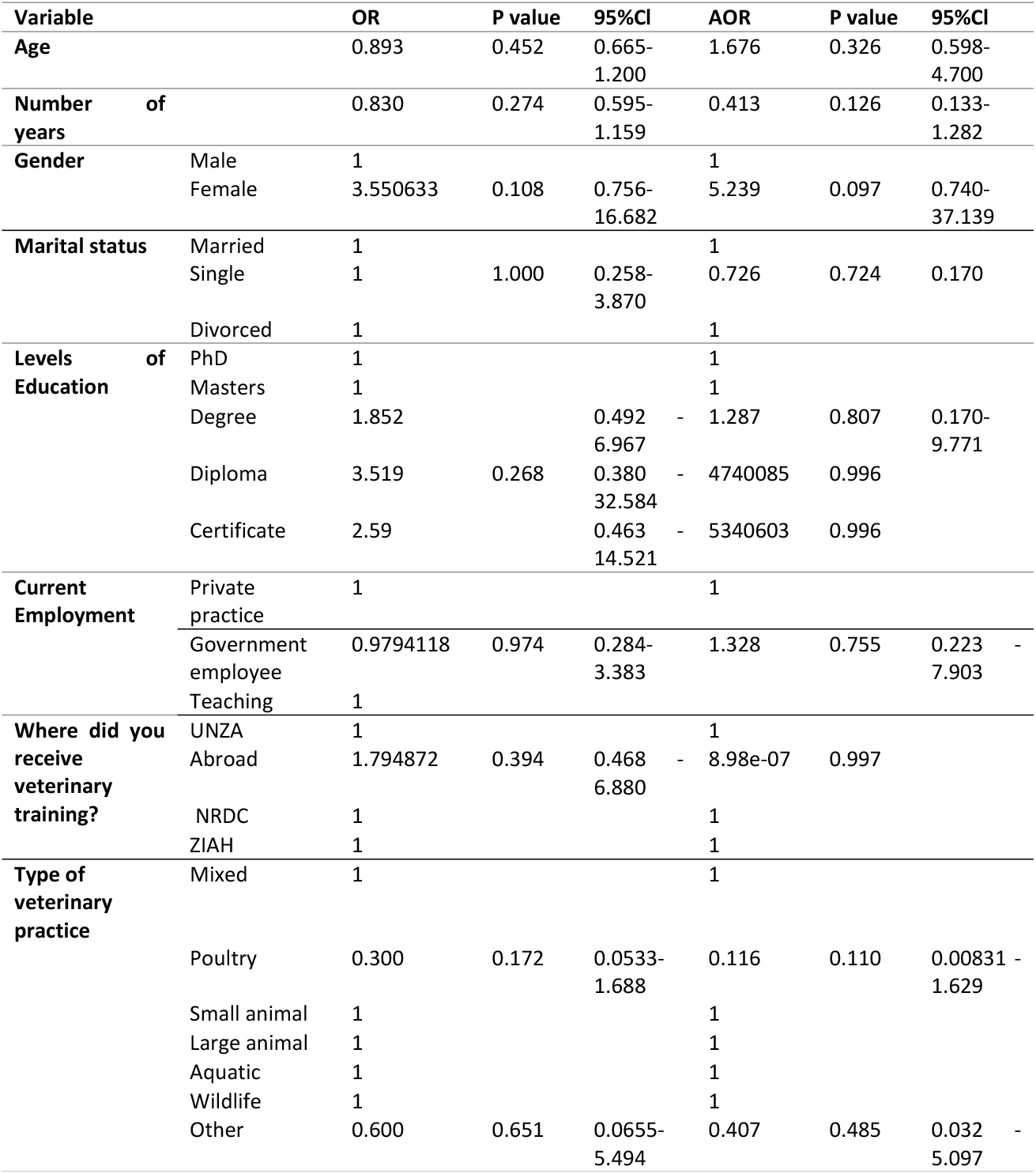
Results of the logistic regression on determinants of knowledge about AMR among Veterinary and para-veterinary professionals.

**Table 6:** Results of the logistic regression on determinants of attitudes on AMR among veterinary and para-veterinary professionals.

| Variable |  | OR | P value | 95%CI | AOR | P value | 95%CI |
| --- | --- | --- | --- | --- | --- | --- | --- |
| Age |  | 0.866 | 0.375 | 0.631-1.189 | 0.964 | 0.949 | 0.316-2.938 |
| Number of years of experience |  | 0.816 | 0.266 | 0.570-1.167 | 0.637 | 0.468 | 0.189-2.149 |
| Gender | Male | 1 |  |  | 1 |  |  |
|  | Female | 6.500 | 0.078 | 0.808-52.293 | 9.184 | 0.092 | 0.694-121.582 |
| Marital status | Married | 1 |  |  | 1 |  |  |
|  | Single | 0.784 | 0.732 | 0.196-3.142 | 0.469 | 0.443 | 0.0677-121.582 |
|  | Divorced | 1 |  |  | 1 |  |  |
| Levels of Education | PhD | 1 |  |  | 1 |  |  |
|  | Masters | 1 |  |  | 1 |  |  |
|  | Degree | 1.034 | 0.965 | 0.230-4.649 | 0.408 | 0.450 | 0.0397-4.184 |
|  | Diploma | 1.966 | 0.571 | 0.190-20.321 | 1197060 | 0.995 |  |
|  | Certificate | 1.448 | 0.697 | 0.225-9.331 | 1391113 | 0.995 |  |
| Current Employment | Private practice | 1 |  |  | 1 |  |  |
|  | Government employee | 0.774 | 0.653 | 0.205-2.696 | 0.931 | 0.937 | 0.156-5.551 |
|  | Teaching | 1 |  |  | 1 |  |  |
| Where did you receive veterinary training? | UNZA | 1 |  |  | 1 |  |  |
|  | Abroad | 1.400 | 0.9846319 | 0.353-5.556 | 1.47e-06 | 0.995 |  |
|  | NRDC | 1 |  |  | 1 |  |  |
|  | ZIAH | 1 |  |  | 1 |  |  |
| Type of Veterinary Practice | Mixed | 1 |  |  | 1 |  |  |
|  | Poultry | 0.267 | 0.137 | 0.0598-1.522 | 0.0598 | 0.061 | 0.00315-1.136 |
|  | Small Animal | 1 |  |  | 1 |  |  |
|  | Large Animal | 1 |  |  | 1 |  |  |
|  | Aquatic | 1 |  |  | 1 |  |  |
|  | Wildlife | 1 |  |  | 1 |  |  |
|  | Other | 1 |  |  | 1 |  |  |

## Discussion

This study assessed the knowledge, attitudes, and practices of antimicrobial resistance among veterinary and para-veterinary professionals in Zambia. It highlights both strengths and challenges in antimicrobial resistance (AMR) management among veterinary and para-veterinary professionals in Zambia. While there is a moderate to high level of awareness regarding AMR, gaps persist in translating this knowledge into appropriate practices, emphasizing the need for targeted interventions.

The results showed that most professionals correctly identified AMR and its causes: 66.7% understood its definition, 82% recognized that overuse reduces antibiotic effectiveness, and 89.6% acknowledged transmission between animals and humans. Similar trends were observed in other studies (6,15,16) where AMR awareness was generally high, particularly regarding causes and consequences. However, despite this awareness, significant gaps remained in actual prescribing behavior. In Nigeria, for example, many veterinary respondents used antibiotics for non-bacterial infections like viral (82.3%) and parasitic (71%) diseases, highlighting a common disconnect between knowledge and practice (6). Conversely, in Haryana, India, AMR knowledge was less robust. Only 21% identified direct zoonotic transmission, and fewer than half recognized environmental routes (17). This suggests regional variation in training and awareness, with countries like Zambia demonstrating stronger foundational knowledge but still needing support to convert that knowledge into consistent, evidence-based clinical decisions.

Zambian professionals expressed strong positive attitudes toward AMR mitigation. A majority supported collaboration between veterinary and human health sectors (70.1%), routine AMR surveillance (93.1%), and farmer education on alternatives to antibiotics (95.8%). These attitudes are consistent with findings from other studies where professionals emphasized the importance of regulations, public education, and responsible use (6,16,17). This positive attitude base is a crucial asset, indicating a fertile ground for implementing new policies and interventions aligned with the global “One Health” approach.

Despite the strong knowledge base and positive attitudes, the study identified critical gaps in actual practice that pose significant risks to effective AMR management. The prevalent reliance on empirical treatment over diagnostic-guided therapy, particularly the low rates of AST (only 20.8% reported performing it regularly), is a significant concern. These issues were mirrored in Nigeria (6), where over-the-counter (OTC) sales in agrovet stores were common, and few practitioners utilized diagnostic labs consistently. While in Bhutan, Wangmo et al. (16) reported similar risks from farmer-driven antibiotic demand, its stricter enforcement and regulatory structure helped mitigate inappropriate access and usage. In this study, the identified barriers to AST use included: lack of laboratory services (57.64%); cost of AST to owners (24.31%); urgent needs of treatment (11.11%); and long waiting times for results (16.67%) were systemic challenges that directly impeded the translation of knowledge and positive attitudes into best practices. India’s Haryana region also faced comparable challenges, including OTC sales, improper drug disposal, and counterfeit products, suggesting that weak regulation and limited diagnostics are universal AMR drivers in low-resource settings (17).

Furthermore, as shown in this study, the concerning and widespread practice of animal owners administering antibiotics without veterinary supervision (75%), coupled with professionals sometimes recommending antibiotics without thorough examination (36.81%), points to a multifaceted problem rooted in both public behavior and professional practice. These practices facilitate misuse, underdosing, and non-adherence to withdrawal periods, as qualitatively highlighted by professionals’ perceptions of farmer-related practices. The logistic regression analysis revealed that none of the socio-demographic variables were significant predictors of AMR knowledge, mirroring findings from Adekanye et al (6). This suggests that a general baseline of fundamental AMR knowledge is widely distributed but also underscores that formal education alone does not guarantee sustained or practical AMR understanding. In contrast, Bhutan and the UK reported stronger associations between experience and knowledge, primarily due to structured continuing professional development (CPD), access to guidelines, and institutional support (15,16). This suggests that ongoing training and practical exposure, rather than academic credentials or tenure, play a more decisive role in AMR competence. These cross-country comparisons reveal a shared pattern: high awareness and positive attitudes existed among veterinary professionals, but structural barriers, including poor diagnostic infrastructure, weak enforcement, and lack of access to CPD or guidelines, impeded effective AMR control.

A notable limitation of this study is the inability to distinguish between veterinary and para-veterinary professionals during analysis. This was mainly due to many para-veterinary professionals upgrading their qualifications, and the survey tool not capturing this detail. As a result, differences in knowledge and attitudes between these groups remain unexplored.

Future studies should separate these groups to provide more detailed insights into training needs and stewardship practices across different professional groups.

Regarding the study’s policy implications, the findings underscore the urgent need for comprehensive policy interventions to address antimicrobial resistance (AMR) within the veterinary sector in Zambia. While the study reveals encouraging levels of knowledge and positive attitudes among veterinary and para-veterinary professionals, it also exposes persistent gaps in practice, mainly driven by systemic limitations such as inadequate access to diagnostic services, economic constraints, and weak regulatory enforcement.

To begin with, there is a pressing need to invest in veterinary diagnostic infrastructure, particularly the expansion and decentralization of antimicrobial sensitivity testing (AST) facilities. The limited availability and accessibility of such services were identified as a significant barrier to responsible antibiotic use. Without reliable diagnostics, professionals are often compelled to rely on empirical treatment, which increases the risk of inappropriate prescribing and the emergence of resistance. Strengthening diagnostic capacity at provincial and district levels would empower practitioners to make evidence-based decisions and reduce unnecessary antibiotic use.

Furthermore, continuing professional development (CPD) on antimicrobial stewardship (AMS) should be institutionalized and made mandatory for all veterinary and para-veterinary professionals. The study’s findings suggest that formal education alone is not a sufficient predictor of good AMS practices. This may be influenced by field-level challenges such as limited access to diagnostics, resource constraints, and client demands that compel professionals to prioritize immediate, practical solutions over recommended guidelines. Ongoing, targeted training, especially on rational drug use, antimicrobial stewardship, and emerging resistance patterns, can ensure that practitioners remain updated with current best practices. Regulatory bodies such as the Veterinary Council of Zambia can play a pivotal role in implementing this recommendation by linking CPD compliance to annual license renewal. Additionally, the formulation and enforcement of national AMR stewardship guidelines tailored to veterinary use are essential. Although professionals showed good knowledge and attitudes toward AMR, the absence of clear, enforceable protocols contributes to variation in practice and undermines stewardship efforts. National guidelines should outline appropriate prescribing behaviours, withdrawal periods, record-keeping, and conditions under which AST should be prioritized. These guidelines must be supported by inspection and enforcement mechanisms to ensure compliance across both public and private sectors.

Public awareness and farmer education are also critical policy areas. The high rate of antibiotic use initiated by animal owners without veterinary supervision highlights the need to educate livestock keepers on the risks of misuse. Integrating AMR awareness into agricultural extension services and veterinary outreach programs can foster responsible use at the community level.

Finally, the study supports the need for a One Health approach that promotes collaboration between animal and human health sectors. While the focus was on veterinary and para-veterinary professionals, the findings revealed challenges such as limited access to diagnostics, weak surveillance systems, and low awareness among farmers. These gaps point to the need for greater cross-sector collaboration. Joint training with the human health professionals, shared diagnostic services, and coordinated community education efforts could enhance the effectiveness of AMR response through a more integrated, sustainable One-Health framework.

## Conclusions

This study reveals that while veterinary and para-veterinary professionals in Zambia possess moderate to high levels of knowledge and positive attitudes toward antimicrobial resistance (AMR), there is a significant disconnect between knowledge and practice. Risky behaviours such as empirical antibiotic use without diagnostics, infrequent antimicrobial sensitivity testing, and unsupervised antibiotic use remain prevalent due to systemic barriers like limited laboratory infrastructure, cost, and lack of enforcement. Addressing AMR in Zambia, therefore, will require a comprehensive approach that combines regulatory enforcement, increased investment in laboratory capacity, and practical, accessible training for both professionals and farmers.

## Data availability statement

The original contributions presented in the study are included in the article.

## Acknowledgments

We are grateful to all practicing veterinarians and para-veterinary professionals for facilitating data collection. We are also grateful to Mr Nicholas Chintu Sande for developing the Map based on the percentage response rate per province. AI was used to improve the language of this article: OpenAI. (2025). ChatGPT [Large language model]. https://chatgpt.com.

